# *In vitro* profiling of endocrine cell death using UCHL1 and GAD65 as soluble biomarkers

**DOI:** 10.1101/076513

**Authors:** Benedicte Brackeva, Sarah Roels, Geert Stangé, Gamze Ates, Olivier R. Costa, Zhidong Ling, Frans K. Gorus, Geert A. Martens

## Abstract

**BACKGROUND:** Pancreatic islet grafts are cultured *in vitro* prior to transplantation and this is associated to a variable degree of beta cell loss. Optimization of culture conditions is currently hampered by the lack of a specific and sensitive *in vitro* indicator of beta cell death.

**METHODS:** We developed a high-sensitivity duplex bead-based immunoassay for two protein-type biomarkers of beta cell destruction, GAD65 and UCHL1, and investigated its proficiency for *in vitro* toxicity profiling on rodent and human beta cells, as compared to a semi-automatic and manual image-based assessment of beta cell death, and *in vivo* after intraportal islet transplantation.

**RESULTS:** Both GAD65 and UCHL1 were discharged by necrotic and apoptotic beta cells proportionate to the number of dead beta cells as counted by microscopic methods. *In vitro*, UCHL1 was superior to GAD65, in terms of biomarker stability providing more sensitive detection of low grade beta cell death. *In vivo*, however, GAD65 was consistently detected after islet transplantation while UCHL1 remained undetectable.

**CONCLUSION:** The use of soluble biomarkers represents a fast, selective and sensitive method for beta cell toxicity profiling *in vitro*. UCHL1 is superior to GAD65 *in vitro* but not *in vivo*.

## Introduction

Intra-portal transplantation of human islets in chronic type 1 diabetes (T1D) patients can improve metabolic control and quality of life [1]. Donor-derived grafts are cultured *in vitro* prior to transplantation under standardized culture conditions i) to allow preferential survival of functional beta cells and deplete the preparations of debris, acinar cells, leukocytes and major histocompatibility complex (MHC)-class II positive cells; ii) to determine the cellular composition of the grafts by immunocytochemistry and electron microscopy; iii) to define functional properties of individual human beta cells (DNA and insulin content, rate of glucose-induced insulin release and biosynthesis) and finally iv) to be able to pool grafts from various donors in order to transplant a sufficient number of insulin secreting cells, e.g. according to our experience minimally 2 × 10^6^ cells per kilogram of body weight [2–4]. Long-term *in vitro* culture of islet grafts is, however, also associated to a variable degree of beta cell loss. Optimization of culture conditions to minimize beta cell loss is currently hampered by the lack of a specific and sensitive indicator of beta cell toxicity that can be used in high-throughput. Here we describe the *in vitro* performance of a newly developed duplex immunoassay for two protein-type indicators of beta cell injury. Specificity for beta cell-selective death was achieved by combining two previously described protein-type biomarkers of beta cell destruction: the classical beta cell auto-antigen 65kDa subunit of glutamate decarboxylase (GAD65), and the neuroendocrine-selective ubiquitin carboxy-terminal hydrolase 1 (UCHL1) [5, 6]. Both proteins are reported to be highly beta cell selective and absent in all exocrine cells (acinar and duct) and other endocrine (alpha) cells. High analytical sensitivity was achieved through the development of a duplex enhanced-sensitivity cytometric bead assay (CBA). Implemented on less sensitive immunoassay platforms, both biomarkers were previously successful for the detection and quantification of massive, synchronous necrotic beta cell destruction *in vitro* and *in vivo* [5, 7]. However, during late-stage islet graft rejection, during the natural course of T1D [8] and during long-term islet graft culture *in vitro*, apoptosis is likely the main form of beta cell death Here we investigated the ability of the newly developed sensitive duplex UCHL1-GAD65 CBA assay in comparison to manual or semi-automated high-content microscopy using vital dyes as gold standard, to detect and quantify – rat beta cell apoptosis induced *in vitro* by a pathophysiologically relevant cocktail of pro-inflammatory cytokines (TNF-α, IL-1β and IFN-γ) [9–11], or after induction of endoplasmic reticulum stress by thapsigargin [9, 12–15]. We found that *in vitro* profiling of UCHL1-GAD65 discharge was not only more sensitive and accurate than microscopy-based methods, but also has a broader application field e.g. the time-resolved monitoring of cell death of beta cells physiologically cultured as tight endocrine aggregates.

## Materials and methods

### Beta cell isolation

Rat beta cells were FACS-purified from islet-cell enriched suspensions [16] prepared from adult (8 or 10-week-old, male) Wistar rats (Janvier Bioservices, France). Beta cell preparations consisted of ≥ 95% endocrine cells (90% insulin+, 3% glucagon+, 1% somatostatin+ and 2% pancreatic polypeptide+ cells) and < 2% exocrine cells. Beta cells were cultured in Ham's F10 media (Thermo Fisher Scientific Inc., USA) supplemented with 2 mmol/L L-glutamine, 50 mmol/L 3-isobutyl-1-methylxanthine, 0.075 g/L penicillin (Sigma Aldrich, USA), 0.1 g/L streptomycin (Sigma Aldrich, USA), 10 mmol/L glucose, and 10 g/L charcoal-treated bovine serum albumin (factor V, radioimmunoassay grade; Sigma Aldrich, USA).

### In vitro toxicity profiling

Isolated single rat beta cells (6000 cells/well in 100 µL medium) were cultured in poly-D-lysine-coated 96-well plates (Thermo Fisher Scientific Inc., USA). After seeding, the cells were exposed (on day 0) for 3 to 6 days to the following toxins: i) thapsigargin (0.2 µM final concentration, Sigma Aldrich, USA); ii) a combination of three cytokines: recombinant TNF-α, recombinant human IL-1β and recombinant IFN-γ (all three used at 1ng/mL final concentration, R&D Systems, USA) and iii) hydrogen peroxide (100mM H_2_O_2_final concentration, Merck KGaA, Germany). Medium was replaced on day 3. At day 3 and day 6 supernatant samples were taken for biomarker analysis with duplex cytometric bead array (CBA) and cell viability was assessed according to following methods.

### Cell viability – automatic versus manual

The percentages of viable, necrotic, and apoptotic cells were assessed in the single cell preparations after a culture period of 3 to 6 days. For this purpose, the cells were exposed to the DNA binding dyes Hoechst 33342 (Ho; Sigma Aldrich, USA) and propidium iodide (PI; Sigma Aldrich, USA)[14]. Hoechst 33342 readily enters cells with intact membranes as well as cells with damaged membranes whereas uptake of the highly polar dye PI is restricted to cells with loss of plasma membrane potential, staining only dead cells. After 15 min of incubation at 37°C with 5% CO_2_, cell viability was assessed both automatically (operator independent) and manually (operator dependent). Absolute number of cells was determined automatically by whole-well imaging using Pathway 435 (BD Biosciences, USA) (Fig 1A). Acquisition of pictures using customer defined assays is done with Attovision software (BD Biosciences, USA). After fine autofocus of Hoechst stained nuclei, a whole well montage of 8 by 12 pictures is taken from the Hoechst channel and from the PI -channel. Pictures are analysed by Attovision software (BD Biosciences, USA); a signal threshold is set on reference pictures made from each pictures by RB25 and “sharpened hat” algorithms. After watershed, upper and lower limits are set to exclude debris and the number of viable and or dead (PI+) cells is calculated for each well.

The cells were manually examined in an inverted fluorescence microscope with ultraviolet excitation wavelength at 340–380 nm. Viable or necrotic cells were identified by intact nuclei with, respectively, blue (Ho) or pink (Ho plus PI) fluorescence. Apoptotic cells were detected by their fragmented nuclei which exhibited either a blue (Ho) or pink (Ho plus PI) fluorescence depending on the stage in the process (Fig 1B). In early apoptosis, only Hoechst 33342 will reach the nuclear material, whilst in a later phase PI will also penetrate the cells. In each condition and experiment, a minimum of 300 cells were counted. Percentages of living, apoptotic, and necrotic cells were expressed as mean ± SD.

## Detection of UCHL1 and GAD65

### Detection antibodies

UCHL1 was detected by a sandwich antibody couple consisting of a capture antibody, polyclonal goat anti-UCHL1 (Acris Antibodies, USA) raised against a peptide corresponding to amino acids 58-68) and rabbit polyclonal anti-UCHL1 detection antibody (Atlas Antibodies, Sigma Aldrich, USA), raised against a peptide corresponding to amino acids 74-219) [5]. GAD65 was detected by previously described antibodies [17]. The detection antibodies were biotinylated using EZ Link™ NHS-Biotin (Thermo Fisher Scientific Inc., USA) according to manufacturer's protocol.

### Cytometric bead array (CBA)

The capture antibodies were covalently attached to CBA functional beads A4 and E7 (BD Biosciences, USA) with sulfosuccinimidyl 4-(N-maleimidomethyl) cyclohexane-1-carboxylate chemistry according to manufacturer's protocol. Functionalized beads were stored shielded from light at 4°C for no more than six months. Recombinant proteins were used to generate standard curves and quality controls were stored in single use aliquots at −80°C providing long-term stability (recombinant human UCHL1, Sigma Aldrich, USA and recombinant human GAD65, Diamyd, Sweden). For a single measurement, 25 µL of sample (recombinant standard material or supernatant sample) was mixed with 25 µL of Assay Diluent (BD Biosciences, USA) and 20µL beads solution at 6000 beads per well in Capture Bead Diluent (BD Biosciences, USA). This was incubated in a U-shaped 96-well predilution plate overnight at 4°C, shaken and shielded from light. Hereafter, 20µL of a 5µg/mL solution of biotinylated detection antibody for UCHL1 and GAD65 was added to all wells without prior washing. After 1 hour incubation at room temperature on the shaking plate, wells were transferred to Multiscreen® filtration plate (Merck KGaA, Germany) and washed 6 times with 150 µl of Wash Buffer (BD Biosciences, USA) using a Millipore Multiscreen (Perkin Elmer, USA) adaptor, followed by addition of 100 µL of Human Enhanced Sensitivity Detection Reagent (Part B) (BD Biosciences, USA). After incubation for one hour at room temperature, shaken and shielded from light, wells were washed 6 times and content was transferred to 1.5 mL microcentrifuge tubes for analysis on FACS Aria cell sorter (BD Biosciences, USA). Single beads were gated based on forward scatter, their intrinsic color code defined by Allophycocyanine (APC)-cyanine (CY7) dye couple after excitation by a 633 nm laser. Clustering parameters for the beads were defined by their forward and sidescatter properties and their APC and APC-Cy7 fluorescence excited by a 640 nm laser. Bound biotinylated detection antibody was quantified by quantification of streptavidin-bound phycoerythrin (PE) fluorescence at 488 nm. Frequency distribution histograms of PE-fluorescence showed a single high peak in all samples and the median PE fluorescence of this peak was selected for quantification with Diva software (BD Biosciences, USA). Analytical limit of detection (LOD, mean of blanks + 3SD) in culture medium was 5.99 pmol/L for UCHL1 and 0.27 pmol/L for GAD65.

### Western Blotting

Antibody dilution used for Western blotting on NuPAGE Novex 10% Bis-Tris Midi Gel (Thermo Fisher Scientific, USA): primary detection with polyclonal rabbit anti-UCHL1 (Aviva Systems Biology, San Diego, USA) at 1:1000 dilution and HRP-linked secondary anti-rabbit antibody (1:1000, GE Healthcare limited, UK). Intensities of bands were quantified by ImageJ software and normalized to beta-actin (rabbit anti-ACTB, 1:1000, Abcam, Cambridge, UK).

### In vitro human islet transplant samples

Human islet cells were cultured for maximally 4 weeks in serum-free Ham's F10 medium and characterized for beta cell number, endocrine purity and viability before selection for clinical transplantation as described [4]. In this study, standardized islet grafts (n=14) sampled at the time of intra-portal transplantation were cultured for 24 hours in 24 well plates with 500 µL serum-free Ham's F10 medium. Subsequently, cells and medium were collected, centrifuged, and supernatant from duplicate samples were combined. Human pancreatic islet graft characteristics are described in Table 2.

### In vivo human islet transplant plasma samples

The UCHL1-GAD65 CBA assay was also applied to measure circulating UCHL1 and GAD65 concentrations after islet transplantation: non-uremic T1D patients typically received intra-portal infusion of at least 2.10^6^ beta cells/kg bodyweight and plasma was sampled at various time points before and up to 24 hours after transplantation. Plasma was collected in K3-EDTA Monovette® tubes (Sarstedt) supplemented with 1% of a 0.12mg/mL solution of aprotinin (Stago) in 0.9% NaCl solution. Storage was done at −80°C after centrifugation for 15 minutes at 1600g.

### Ethics Statement

All experiments involving humans and animal models were approved by the medical ethics committee of Universitair Ziekenhuis Brussel and Vrije Universiteit Brussel (Commissie Medische Ethiek, O.G. 016, Reflectiegroep Biomedische Ethiek, Laarbeeklaan 101, B1090 Brussels, Belgium). Use of animal and human tissues for in vitro biomarker validation was covered by B.U.N. 143201213515 for a project entitled “Identification and clinical validation of biological markers for beta cell death or dysfunction”, approval to G. Martens. Use of rat beta cells was covered by project reference 12-274-2 entitled “Use of rats for preclinical transplant experiments”, approval to Prof. Dr. D. Pipeleers. Use of plasma samples obtained from human islet grafts recipients was covered by project reference 2005/136 prot, entitled “Long-term function of pancreatic beta cell allografts in non-uremic type 1 diabetic patients”. Use of human beta cells for in vitro research was covered by project reference CME 2005/118 and CME 2010/193 (approval to the UZ Brussel BetaCell Bank, B.U.N. 14320109289) and by project reference CME 2009/018, entitled “platform for beta cell therapeutics” and CME 2009/017, entitled CME 2009/017, approval to the JDRF Center for Beta Cell Therapy at Vrije Universiteit Brussel.

Human islet grafts that did not meet the criteria for clinical transplantation were provided for secondary in vitro use according to the Belgian law (December 19, 2008 – Wet inzake het verkrijgen en het gebruik van menselijk lichaamsmateriaal met het oog op geneeskundige toepassing op de mens of the wetenschappelijk onderzoek) along the principle of presumed consent, unless an express objection was made by the person or next of kin prior to the death. All organs are traceable through written forms (Eurotransplant Pancreas Report, Organ inspection record, donor feedback record, Beta Cell Graft Release Record, Request forms for human pancreatic tissue/cells) archived at the BetaCell Bank at UZ Brussel.

All human islet graft recipients provided written informed consent to use blood samples, along predefined blood sampling algorithms reviewed by our medical ethics committee. Informed consent forms were made up in Dutch and French, and approved by the medical ethics committee of Universitair Ziekenhuis Brussel and Vrije Universiteit Brussel, and consent forms archived by the UZ Brussel Clinical Trial Center.

### Statistical analysis

Data were expressed as mean ± SD and p-values were calculated with Mann-Whitney U-test using GraphPad Prism (GraphPad Software, USA). Statistical significance is denoted by *p<0.05 and **p<0.0001.

## Results

### Imaging-based measurement of rat beta cell death in vitro

On day 0, 6000 single rat beta cells were seeded and exposed to two known inducers of apoptotic beta cell death [9], ER-stress inducer thapsigargin (0.2 µM) and a cocktail of pro-inflammatory cytokines: TNF-α+IFN-γ (all at 1ng/mL final concentration). Hydrogen peroxide (H_2_O_2_; 100µM final concentration), a potent oxidative inducer of necrotic rat beta cell death was used as positive control [18]. Cell death and viability were measured after 3 and 6 days using an operator-dependent (manual) or -independent (semi-automated) microscopic method. In the semi-automated method, a whole well image was generated using a BD Pathway 435 platform as shown in Fig 1A, the total number of cells was counted by integrating Hoechst-positive events and the number of dead, propidium-iodide positive cells was calculated using BD Attovision software. These same wells were then subjected to manual counting of percentage necrotic, apoptotic and living cells as indicated in Fig 1B and 1C. H_2_O_2_ rapidly induced massive necrosis resulting in > 90% necrotic cells at day 3. On day 3, the thapsigargin -and cytokine-toxicities became detectable with 36.0 (37.3) % and 45.9 (41.9) % dead cells, respectively as detected by the semi-automated (manual) counting. However, this effect failed to reach statistical significance as compared to control cells 35.0 (29.2)%. On day 6, thapsigargin and cytokines resulted in death of 82.9 (88.7) % and 67.1 (63.7) % cells, respectively, and expectedly, the predominant modality of cell death was apoptosis (Fig 1E). The side-by-side comparison between automatic and manual counting of cell death indicated overall an excellent agreement (R^2^ =0.956, Fig 1F) between semi-automated and manual methods, thus providing the required reference to investigate the added value of measuring discharged biomarkers as sensitive indicators of *in vitro* beta cell toxicity.

### Biomarker-based measurement of rat beta cell death in vitro

Loss of beta cell plasma membrane integrity results in the discharge of obligate intracellular proteins that, when sufficiently beta cell-selective, qualify as real-time indicators of this death process. Previous studies, using relatively insensitive immunoassays, indicated that GAD65 and UCHL1 are extracellularly discharged under conditions of severe necrotic destruction [5, 6]. Here, we investigated if a newly developed enhanced-sensitivity bead-based duplex assay for GAD65 and UCHL1 could also detect minor instances of apoptotic beta cell death *in vitro*, taking the imaging-based methods (Fig. 1) as reference. UCHL1 came out as best indicator: (i) Induced UCHL1 discharge by thapsigargin and cytokine-induced cell death was consistently (P < 0.05) detected starting day 3. At this time point, GAD65 detected cytokine-but not thapsigargin-induced cell death (Fig. 2A); (ii) Stoichiometrically, extracellular UCHL1 was 20 times more abundant than GAD65; (iii) the molar amount of UCHL1 discharged between day 3 and 6 showed an overall good agreement with the imaging-based increment of cell death during this time frame (Fig. 2B), while this was not the case for GAD65; (iv) UCHL1, was discharged early during the incubation, as occurs during H_2_O_2_-induced necrosis. It remained stable throughout the full 6 day incubation at 37°C while GAD65 showed 30 % degradation between day 3 and day 6. This is in line with its reported thermolability [19], and was also confirmed in a stability experiment in which GAD65 was spiked in culture medium in absence of cells and maintained at 37°C for 3 days; this resulted in only 19% recovery (Supplementary Fig 1). Overall, the discharged amounts of both UCHL1 and GAD65 showed a significant (P<0.0001) linear correlation to the number of dead beta cells, but expectedly this correlation was the strongest for UCHL1 (Fig. 2C).

**Fig 1.**
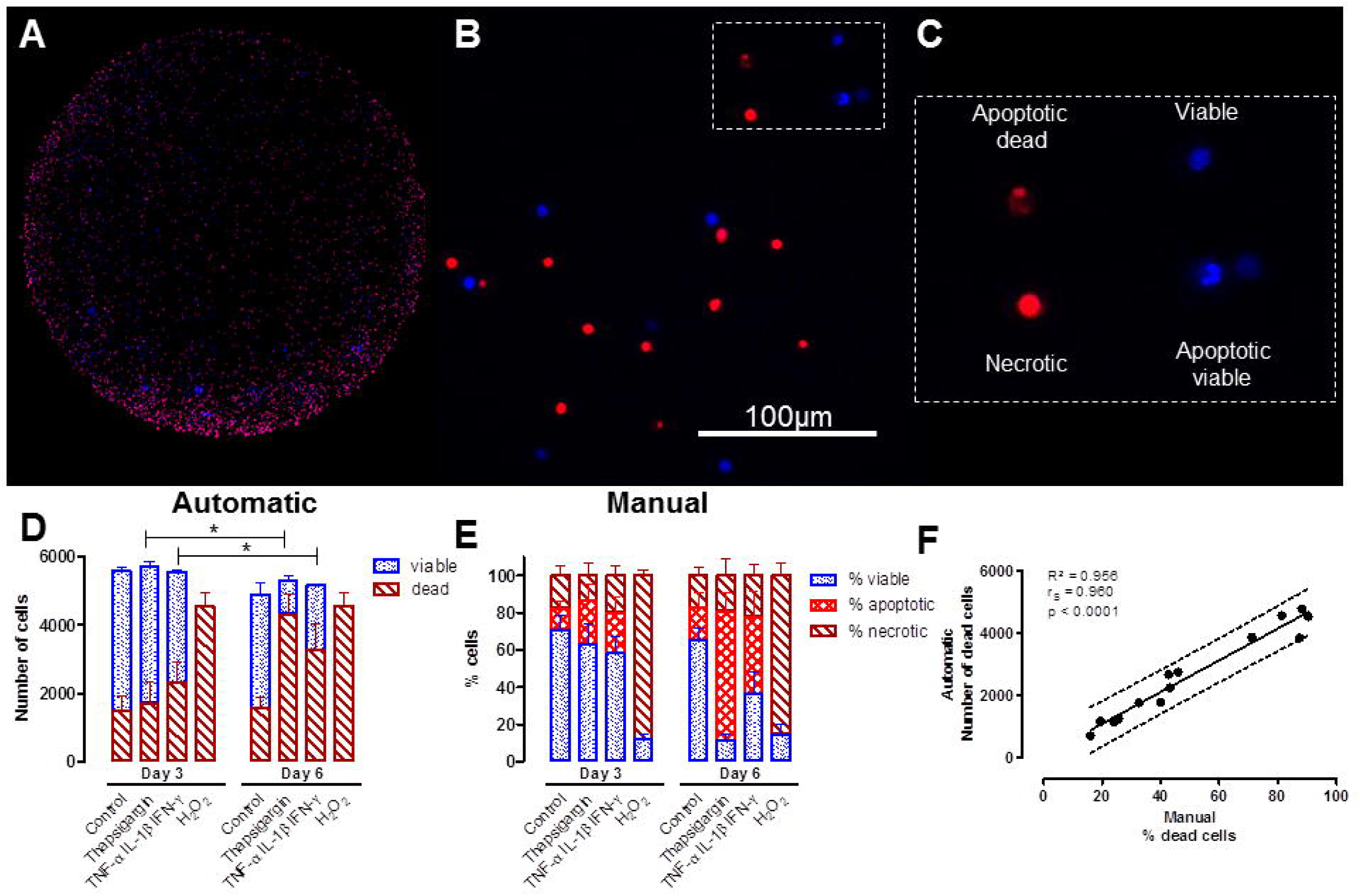
Imaging-based measurement of rat beta cell death *in vitro*. Representative Hoechst-Propidium iodide vital staining of single rat beta cells following exposure to TNF-α+ IL-1β IFN-γ for 6 days. (A) Whole-well image on BD-Pathway 435, segmentation using Image Processing Lab software. Hoechst positive cells in blue and propidium iodide positive cells in red. (B) Single rat beta cells, representative image for manual counting. Scale bar indicates 100µm. (C) Inset shows detail of panel B with viable (blue), necrotic (red) and live (pyknotic nucleus, blue) and dead (fragmented nucleus, red) apoptotic cells (D) Number of viable/dead cells counted automatically with Attovision software for the different conditions over time (day 3 and day 6). Bars represent mean ± SD (n=2). (E) Percentage of cells accounted to viable, necrotic or apoptotic with manual counting for the different conditions over time (day 3 and day 6). Bars represent mean ± SD (n=2-4). (F) Correlation between number of dead cells counted automatically with Attovision software and percentage of dead cells (necrotic and apoptotic) by manual counting. A positive correlation was found with Spearman r = 0.960, R² = 956, P<0.0001. Dotted lines indicate 95% prediction interval.

**Fig 2.**
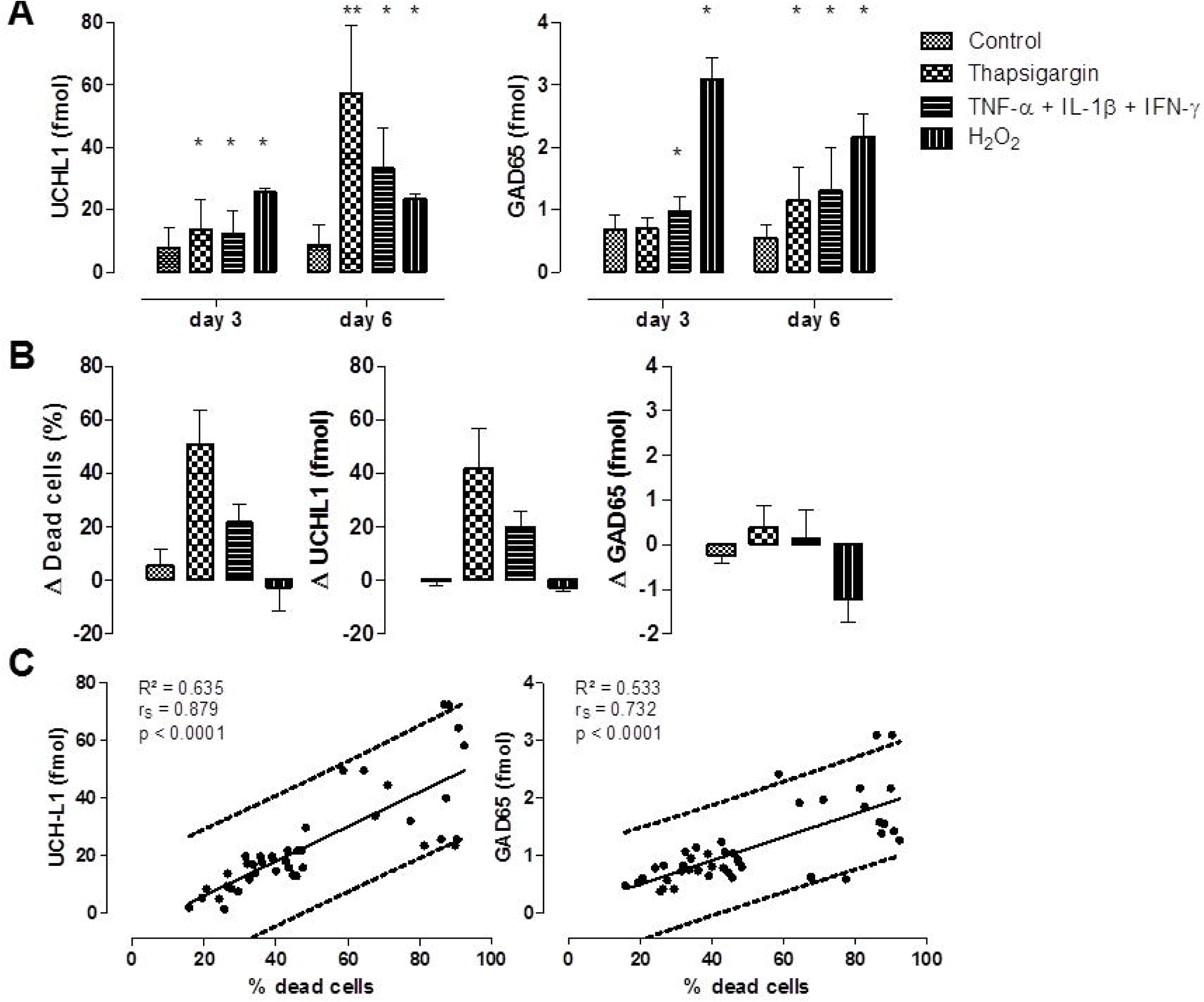
Biomarker-based measurement of rat beta cell death in vitro. (A) UCHL1 and GAD65 protein discharged from beta cells following exposure to thapsigargin (0.2µM), cytokines TNF-α + IL-1β + IFN-α(all 1 ng/mL) or hydrogen peroxide H_2_O_2_(100µM) compared to control (culture medium) at day 3 and day 6. * p<0.05 versus control, Mann Whitney test (B) Difference in cell death (%), UCHL1 (fmol) and GAD65 (fmol) between day 6 and day 3 expressed as delta values. (C) Correlation between UCHL1 and GAD65 (fmol) and % dead β cells as found by manual counting. A positive correlation was found for both proteins with Spearman r = 0,912, P<0.0001 and Spearman r = 0.771, P<0.0001 respectively.

Analysis of biomarker recovery revealed, however, one anomaly for UCHL1: since discharged UCHL1 is extracellularly stable (Fig. S1 and Fig. 2), the amount of released UCHL1 should be equivalent to the intracellular content of the disrupted beta cells. This was not the case: for each K dead beta cell, 9.5 fmol UCHL1 was recovered from the medium, well above the intracellular content measured by the same CBA-method in isolated cells (Table 1). This excessive recovery of UCHL1 *in vitro* was previously observed after severe necrosis by hydrogen peroxide [5]. It could not be explained by increased adaptive UCHL1 biosynthesis under conditions of ER-stress (S2 Fig), leaving the option that it may results from an altered immune reactivity of UCHL1 during extracellular discharge, e.g. by loss of intracellular polymer structure [20]. This might explain higher observed intracellular UCHL1 levels by denaturing methods such as Western blotting (Table 1).

**Table 1.** Intracellular content of UCHL1 and GAD65

| Table 1. Intracellular content of UCHL1 and GAD65 |  |  |  |  |
| --- | --- | --- | --- | --- |
| UCHL1 | RAT |  | HUMAN |  |
| fmol/K |  | n |  | n |
| CBA | 1.1 (0.4-1.3) | 4 | 0.8 (0.7-1.1) | 5 |
| Western Blot | 2.0 (1.2-4.3) | 4 | 9.5 (4.5-18.5) | 8 |
| GAD65 | RAT |  | HUMAN |  |
| fmol/K |  | n |  | n |
| CBA | 3.0 (1.7-4.3) | 3 | 1.1 (0.7-1.6) | 4 |
| Data represent median with IQR in parentheses |  |  |  |  |

### Biomarker-based measurement of human beta cell death in vitro

Human beta cells, prepared for clinical transplantation, survive and function optimally when cultured as endocrine aggregates, referred to as islet equivalents. Since aggregate suspension cultures technically impede image-based measurement of beta cell death, biomarker-based measurements represent a useful alternative. Therefore, we measured UCHL1 and GAD65 discharge *in vitro* over 24h by standardized islet grafts (n=14) sampled at the time of intra-portal transplantation, and studied statistical correlations to the graft characteristics shown in Table 2. The grafts showed a good overall viability and a low level of contamination with exocrine acinar cells, as measured by electron microscopy, and state-of-the-art endocrine purity (95%CI) with 31% (19-35)% insulin-positive cells. The amount of UCHL1 discharged *in vitro* correlated significantly both to the total percentage of dead cells, and to the number of beta cells. Also for GAD65, a correlation could be found with the number of beta cells, but this correlation was less strong compared to UCHL1 and was also seen with total cell number (Table 2).

**Table 2.**
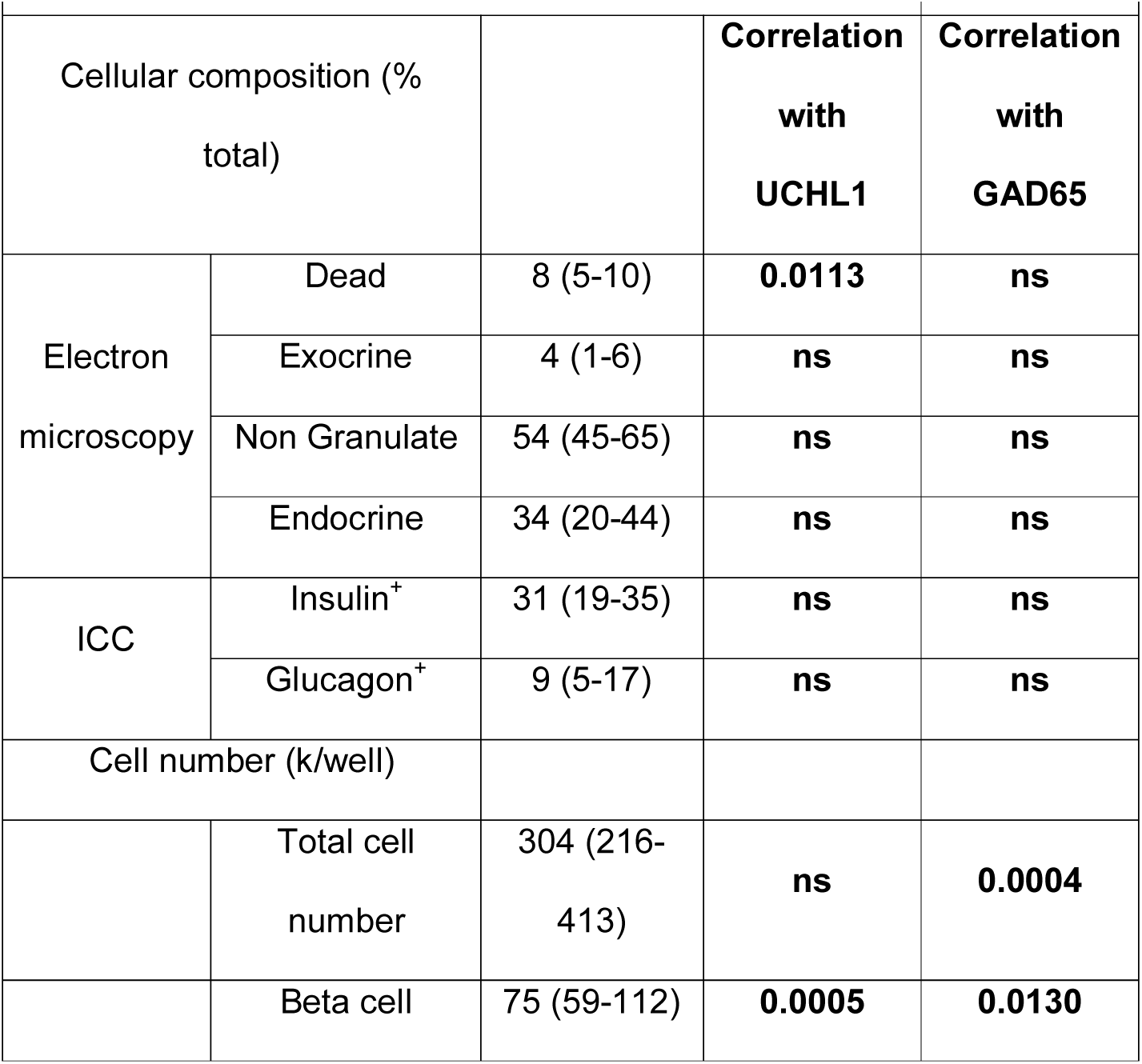
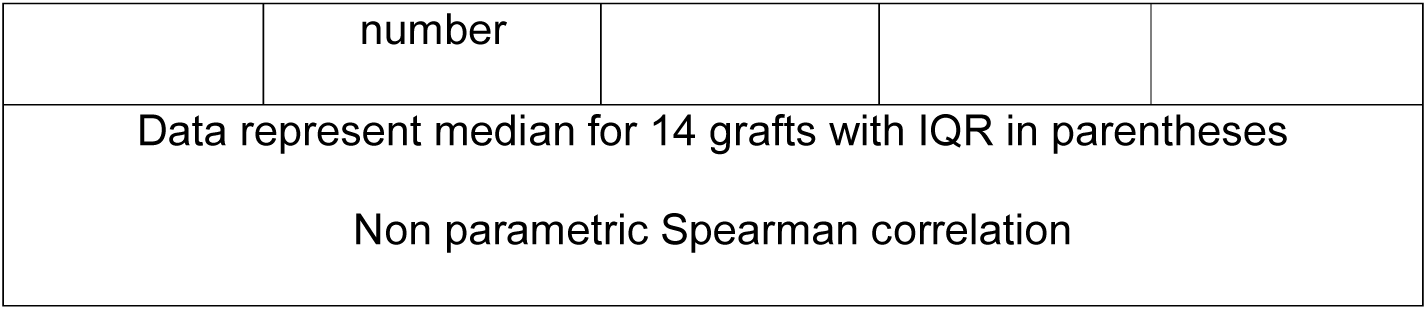
Human pancreatic graft characteristics

### Failure of UCHL1 to detect human beta cell death in vivo

GAD65 was previously validated as real-time indicator of human beta cell death after islet transplantation [21]. UCHL1 could be confirmed as *in vivo* indicator of beta cell death in streptozotocin-injected rodents, albeit variably and hampered by fast hepatic biomarker clearance [5]. We tested the duplex UCHL1-GAD65 CBA assay in GAD65-autoantibody negative islet graft recipients: while an acute plasma GAD65 surge could consistently be detected, UCHL1 remained undetectable, both after the standard implantation in the liver (Fig 3), as after experimental implantation into the omentum, indicating that the superior *in vitro* performance of UCHL1 as compared to GAD65 is not translated into clinical diagnostic use.

**Fig 3.**
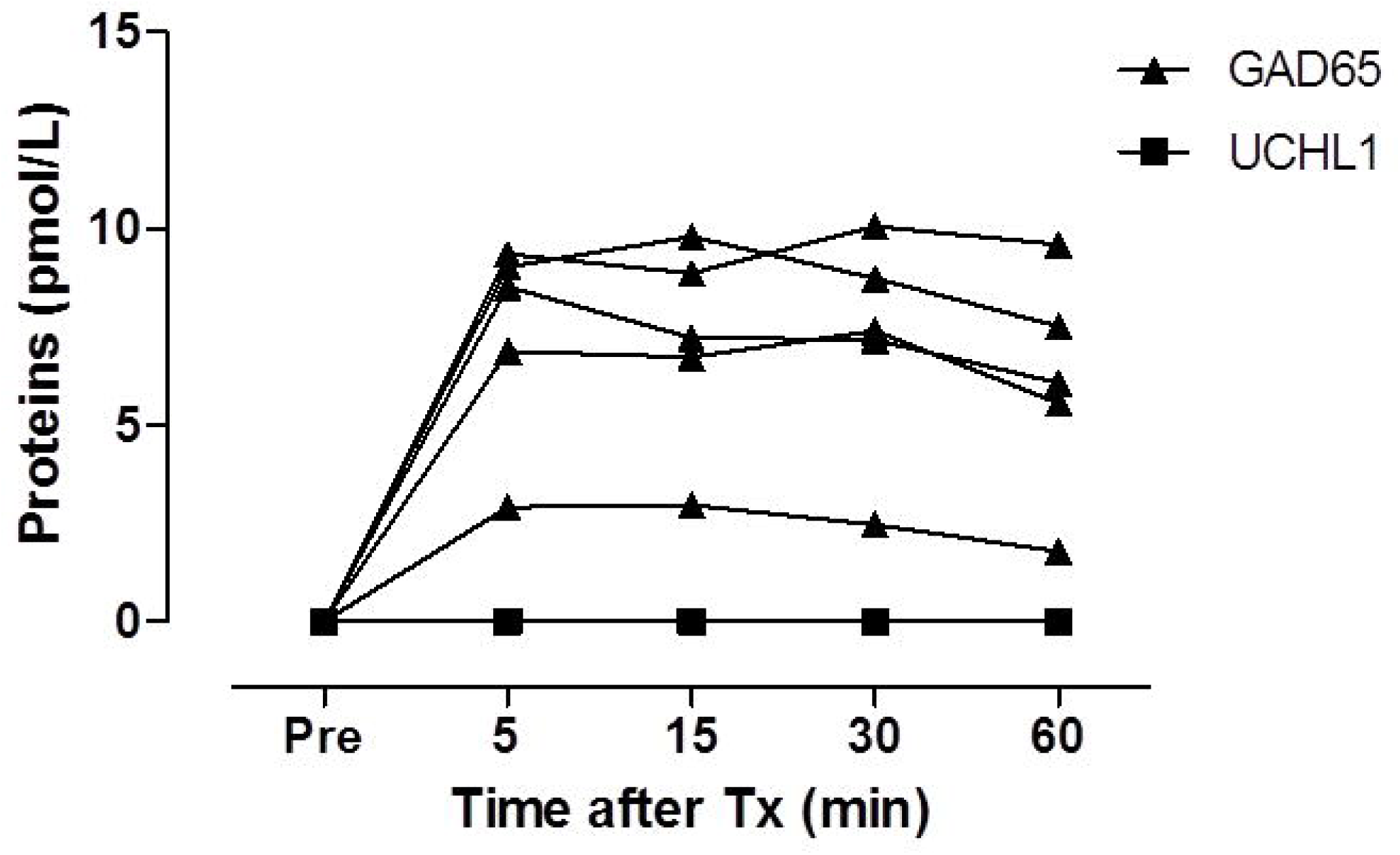
Biomarker-based measurement of human beta cell death *in vivo*. Plasma samples from islet transplantations (Tx) (n=5) were analyzed for their GAD65 (▲ and UCHL1 (■) concentration. Venous blood samples were taken before Tx (Pre) and early after Tx at 5-15-30-60 minutes.

## Discussion and Conclusions

Here we report a novel method for *in vitro* analysis of rodent and human beta cell toxicity using a sensitive duplex UCHL1-GAD65 CBA assay. The quantification of discharged beta cell-biomarkers has several technical advantages for *in vitro* toxicological analysis of beta cells, as compared to microscopic quantification of vital dyes or conventional tetrazolium dye-based (MTT) assays: (i) by targeting endocrine-restricted biomarkers, it can specifically quantify beta cell injury in mixed cultures or impure islet fractions. It performs equally well in adherent and suspension cultures, and unlike MTT, independently of the metabolic phenotype of the cells; (ii) by using an enhanced-sensitivity bead-based immunoassay approach [17], it achieves sensitivities that could, in our hands, not be reached by MTT assays on the small numbers of cells used in our experimental design, achieving even superior sensitivity than the microscopic measurement of vital dyes; (iii) moreover, soluble markers allow repeated sampling of the same wells and, unlike MTT and microscopy, does not represent endpoint studies, thus allowing kinetic time-resolved analyses, and interim sampling for quality control purposes, and finally (iv) the assay allows upscaling to high-throughput formats on widely available immunoassay platforms.

*In vitro*, UCHL1 appears superior to GAD65 as indicator of beta cell injury: UCHL1 showed a better thermo-stability and outperformed GAD65 to detect early stages of beta cell apoptosis, although its unexplained excessive immuno-recovery still necessitates the use of internal controls. We repeatedly observed a 3-7 fold higher recovery of immuno-detectable UCHL1 than could be expected based on measurement of intracellular content. We hypothesize that this behavior is explained by the loss of UCHL1 multimeric structure [23] after extracellular discharge, inducing a higher accessibility of the UCHL1 epitopes targeted by our sandwich immunoassay on the UCHL1 monomers. This behavior requires specific attention, since UCHL1 is increasingly used in patients to quantify neuronal damage [23, 24]. UCHL1 is reported as one of the most abundant proteins in neurons [23]. In human beta cells it ranks among the higher-abundance core proteome detectable by LC-tandem MS [5, 25, 26]. Its neuroendocrine-selectivity, combined with its high cellular abundance, favor the use of UCHL1 as biomarker to detect neuronal or neuroendocrine cell death *in vivo*, Several groups reported that intact UCHL1 protein can pass the blood brain barrier and become detectable in the circulation after acute traumatic or ischemic brain injury [24]. Using our sandwich immunoassay, we could detect beta cell-derived UCHL1 in the plasma after streptozotocin-induced synchronous and massive induction of beta cell necrosis in rats [5]. Here, we show that despite its excellent *in vitro* behavior, the use of UCHL1 failed the translation to the transplantation clinics. In human islet graft recipients, UCHL1 could not be detected after intra-hepatic nor after intra-omental transplantation, although we expected detectable levels as estimated by extrapolation of its *in vitro* discharge compared to GAD65. We hypothesize that the failure of UCHL1 might be attributed to its fast hepatic clearance [5] or, loss of its immune recognition due to polymerization or physiological binding of ubiquitinated-proteins in its catalytic center. If the latter is true, this would also predict variability of immuno-detectable UCHL1 in other models (e.g. traumatic brain injury), and caution in the use of UCHL1 as quantitative indicator of neuroendocrine cell death.

In conclusion: the use of soluble biomarkers represents a fast, selective and sensitive method for beta cell toxicity profiling *in vitro*. UCHL1 is superior to GAD65 *in vitro* but not *in vivo*.

## Acknowledgements

This study was supported by research grants from the Research Foundation Flanders (FWO G.0492.12 project grant and Senior Clinical Investigator career support grant to G.M.), from the European Foundation for the Study of Diabetes (EFSD/JDRF/Lilly Programme 2015 Award to G.M. and F.G.), from the JDRF through an Innovative Pilot Grant (Pilot Studies Relevant to Biomarkers of Human Type 1 Diabetes, 1-PNF-2014-181-A-V), by the Agentschap voor Innovatie door Wetenschap en Technologie (IWT to G.M. and O.C.) and by the Wetenschappelijk Fonds Willy Gepts from the Universitair Ziekenhuis Brussel (to G.M). Funding organizations did not influence data collection and interpretation. We thank Prof. Dr. Daniel Pipeleers for his scientific and logistic support.

## Supplementary figure legends

**Supplementary Fig. 1:**
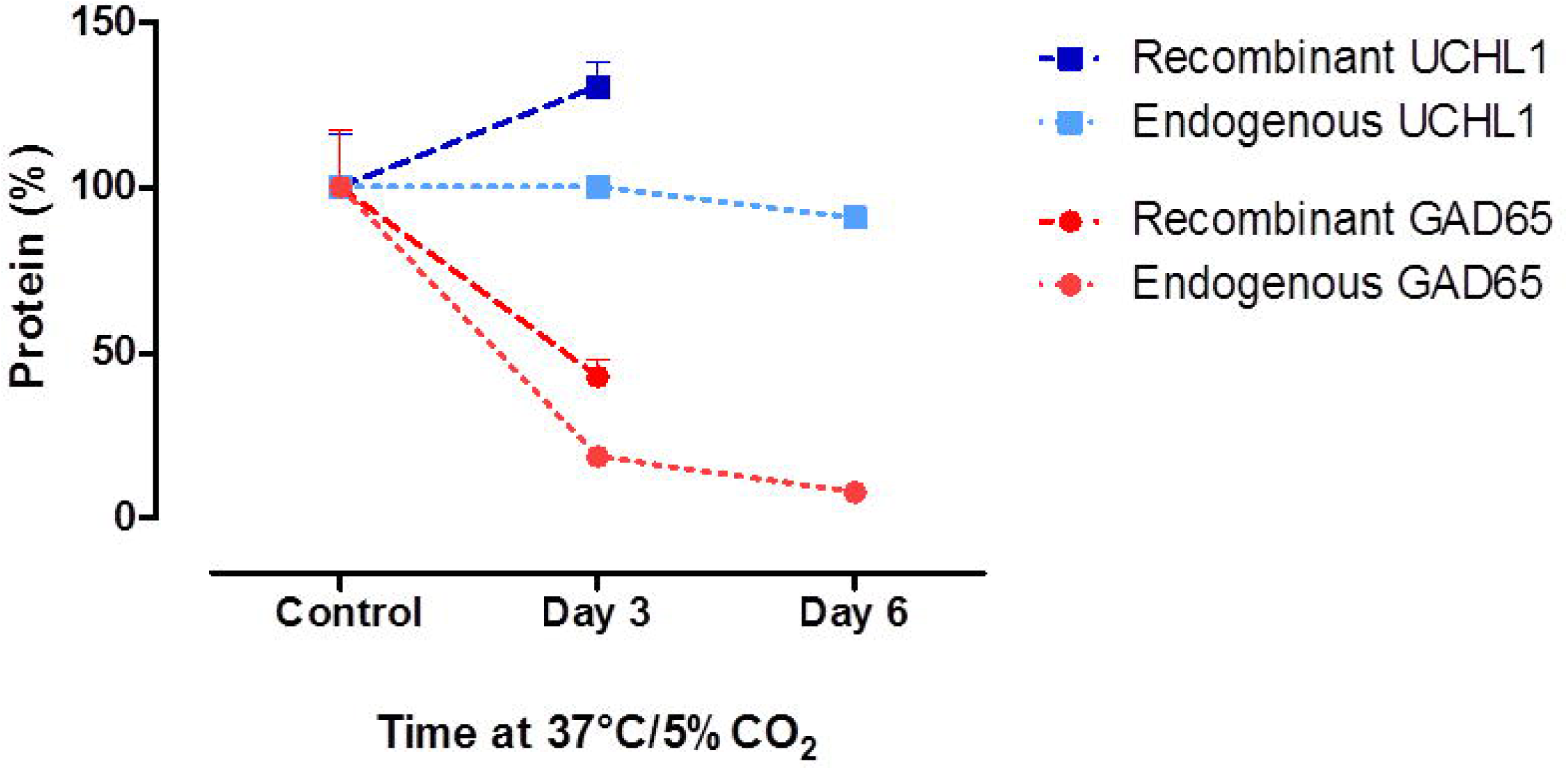
Stability of recombinant and endogenous proteins. UCHL1 (blue) and GAD65 (red) spiked in culture medium and stored at 37°C and 5%CO_2_ for 3 to 6 days.

**Supplementary Fig 2.**
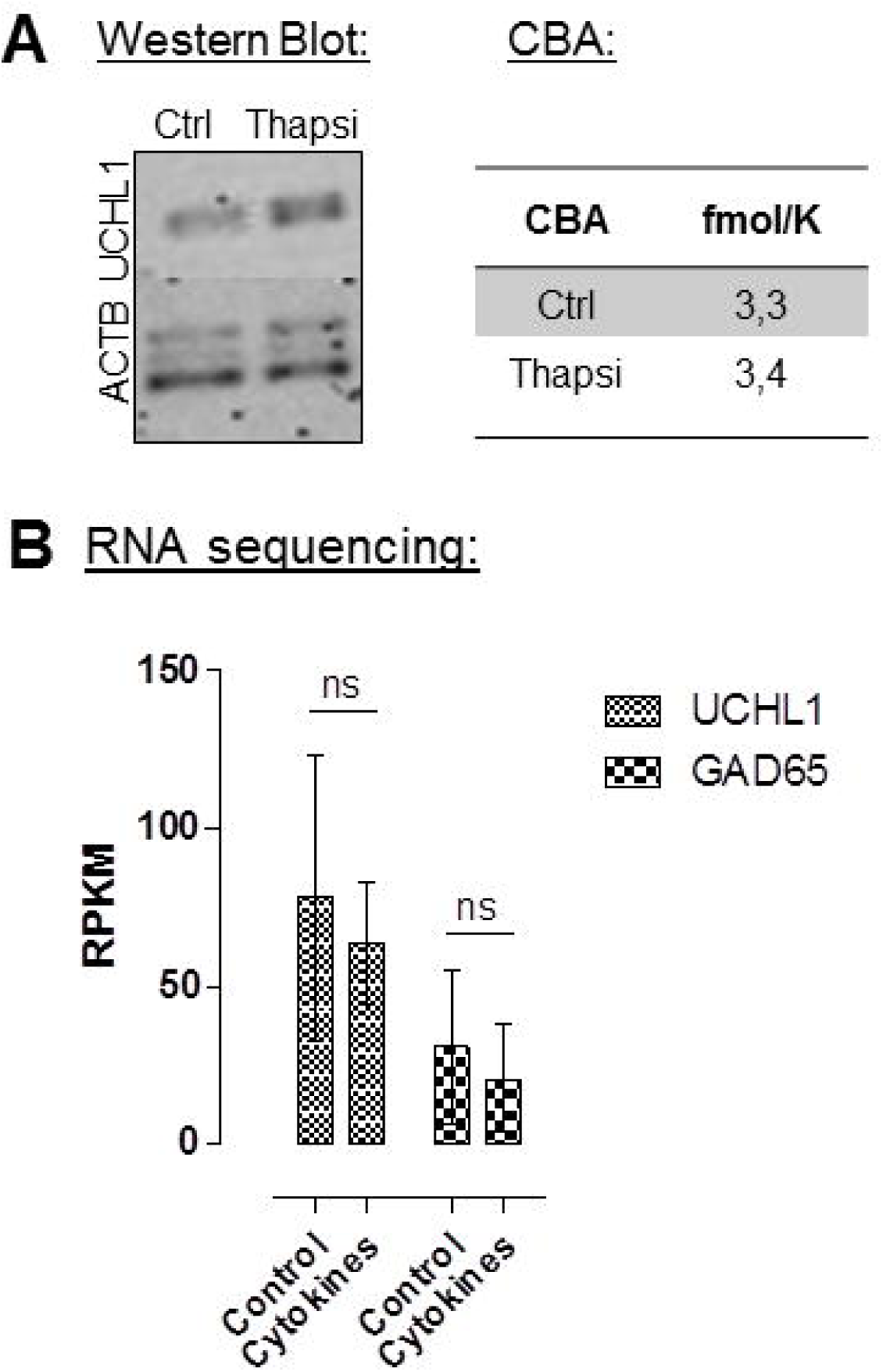
No adaptive UCHL1 biosynthesis under conditions of ER-stress. Rat beta cell aggregates were exposed to thapsigargin (0.2µM) for 24 hours. Cell lysate was compared to control condition (aggregates exposed to culture medium) and analysed with Western Blotting and with duplex UCHL1-GAD65 cytometric bead assay (A). Both analysis revealed no significant difference between rat aggregates exposed to stress and control condition. (B) UCHL1 and GAD65 mRNA data from previously published dataset [27]. RNA sequencing was used to identify transcripts, including splice variants, expressed in human islets under control conditions or following exposure to the pro-inflammatory cytokines IL-1β and IFN-γ. This revealed no significant difference between human islets exposed to cytokines or control condition.

